# A Novel Denitrification Regulator, Adr, Mediates Denitrification Under Aerobic Conditions in *Sinorhizobium meliloti*

**DOI:** 10.1101/437079

**Authors:** Coreen M. Slape, Arati V. Patankar, Juan E. GonzáLez

## Abstract

*Sinorhizobium meliloti* is a soil dwelling bacteria capable of forming a symbiotic relationship with several legume hosts. Once symbiosis is established, *S. meliloti* fixes atmospheric nitrogen into nitrogenated compounds, thus carrying out an important step in the nitrogen cycle. *S. meliloti* is also capable of the reverse process, denitrification, the reduction of nitrate and nitrite to nitrogen gas. In this study we have identified a novel regulator of denitrification in *S. meliloti*, Adr, which affects the expression of the denitrification genes in aerobically grown cultures. Analysis of the Adr sequence reveals a LuxR-like quorum sensing regulator, however, it does not respond to the known quorum sensing signals produced by *S. meliloti*. Additionally, we show that FixJ, the major regulator of denitrification and microaerobic respiration in *S. meliloti*, is active under our growth conditions. Comparison of the FixJ microarray to our Adr microarray shows a significant overlap between the two regulons. We also show that while Adr is not necessary for symbiotic nitrogen fixation, a functional copy of this regulator confers a competitive advantage to *S. meliloti* during host invasion. Our findings suggest that Adr is a new type of denitrification regulator and that it acts at the same regulatory level as FixJ.

**Importance:** Rhizobia contribute to the nitrogen cycle by fixing atmospheric nitrogen to nitrogenated compounds and by denitrification, the reduction nitrate and nitrite to nitrogen gas. Denitrification enhances the survival of *Sinorhizobium meliloti* in the various environments it may encounter, such as free-living conditions in the rhizosphere, during invasion of the plant host, and after a symbiotic relationship has been established. Oxygen concentration is the typical signal for denitrification gene expression. Recent studies of low oxygen cultures of *S. meliloti* have outlined the regulation structure for denitrification. In this study, we examine the regulation of denitrification in aerobically grown *S. meliloti* cultures. Understanding how *S. meliloti* responds to various oxygen concentrations will result in a more complete picture of denitrification regulation in this agriculturally important organism and the impact of denitrification on the soil microbiome as a whole.

## Introduction

*Sinorhizobium meliloti* is an aerobic soil dwelling α-proteobacteria that is found in a variety of environments in which oxygen concentrations fluctuate, including free-living in the soil or in association with a plant host. Like many *Rhizobiaceae, S. meliloti* contributes to the nitrogen cycle, either by fixing atmospheric nitrogen as bacteroids in conjunction with a legume host (*Medicago sativa*) or by breaking down nitrates and nitrites to nitric oxide, nitrous oxide, or dinitrogen gas (Figure 1) (1). Although these two processes seem incongruous, both play an important role in *S. meliloti* (2, 3). During symbiosis with a legume host, oxygen limitation is intrinsic to the nodule and is required for the expression and function of the nitrogenase enzyme. The first step of denitrification, the reduction of nitrate to nitrite, removes any excess reducing power that may be present in the cell (4). Further steps remove nitrite from the nodule, ensuring the optimal environment for nitrogenase function. Aside from the presence of NO_x_ (NO_3_^−^, NO_2_^−^, NO, or N_2_O), the major signal for the expression of the denitrification pathway is oxygen limitation. Denitrification usually occurs under anaerobic or low oxygen tension conditions where oxygen cannot serve as an efficient electron acceptor. However, it is now well established that denitrification can occur aerobically, either with no or partial oxygen limitation, in several bacteria, including *Paracoccus denitrificans*, *Pseudomonas aeruginosa,* and *Agrobacterium sp*. (5). While denitrification is documented in *S. meliloti,* most reports focus on microaerobic conditions such as those found in the nodule (6, 7). The primary role of aerobic denitrification is likely the removal of excess reducing power or detoxification of NO_x_ found in the environment (5). Since respiratory reduction of NO_x_ is coupled with energy generation, denitrification even in the presence of oxygen can enhance bacterial survival in environments where the oxygen concentration may fluctuate (5).

**Figure 1.**
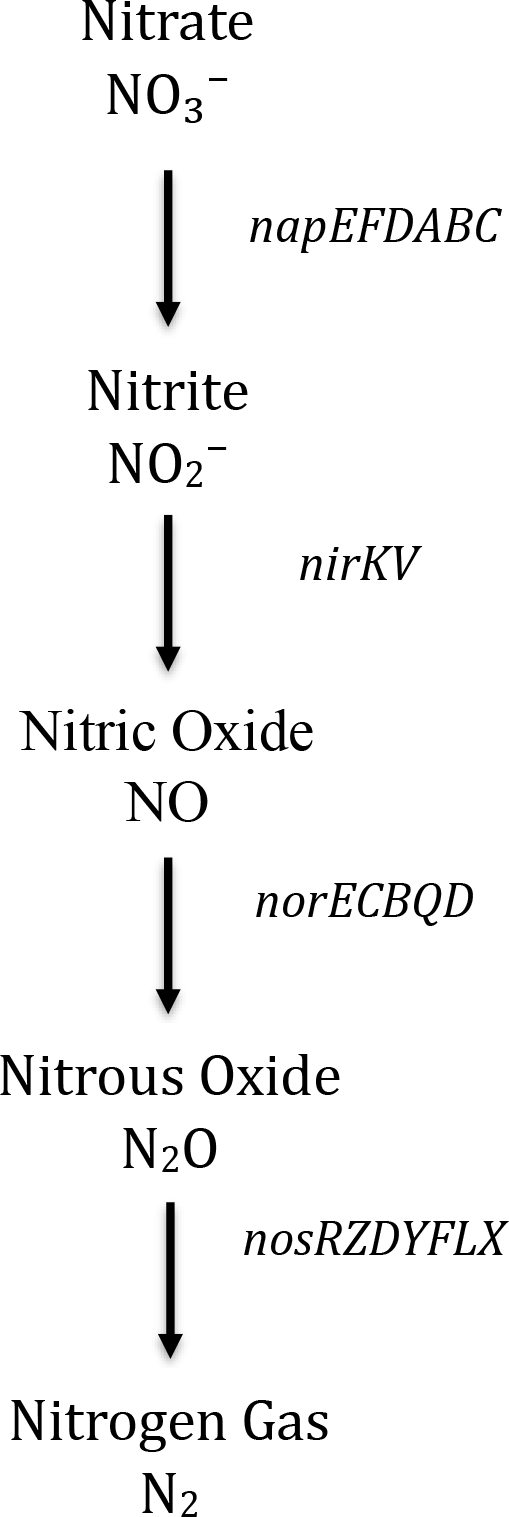
The denitrification pathway in *S. meliloti* and its associated genes.

Denitrification may also allow *S. meliloti* to remove nitrate or nitrite from the soil near plant roots as these compounds inhibit bacterial attachment to host roots (8). While migrating through the infection thread during root invasion, the ability of *S. meliloti* to denitrify enhances its survival when it encounters plant defense responses such as nitric oxide bursts (9). Once *S. meliloti* successfully inhabits the nodules, denitrification is thought to be active to reduce nitrate, as it is inhibitory to nitrogenase activity (10).

An overview of the regulation of denitrification and nitrogen fixation in *S. meliloti* is shown in Figure 8. The FixL/FixJ two component system is essential for *S. meliloti* to form a symbiotic relationship with a legume host (11). FixL is a membrane bound sensory kinase that detects oxygen levels in the environment. When the oxygen concentration in the environment drops, FixL autophosphorylates and transfers the phosphate to FixJ (12). Previous studies have demonstrated that phosphorylation of FixJ relieves the weak interaction of the FixJ DNA binding domain (C-terminal) and the signal recognition domain (N-terminal) by triggering a conformational change (13). Once this interaction between domains is abolished, FixJ is able to bind DNA and/or dimerize (14). The FixJ C-terminal domain alone is able to recognize promoters and activate transcription in the absence of phosphorylation, which supports the theory that the N-terminal signal recognition domain is regulatory in nature (15). Though dimerization is not essential for DNA binding or promoter recognition, it is thought to contribute to promoter binding affinity (14).

Two component systems similar to FixL/FixJ are fairly ubiquitous in nitrogen fixing bacteria. However, regulation downstream of FixJ can vary greatly (16). In *S. meliloti* FixJ has five direct targets: *nifA*, *fixK1*, *fixK2*, *proB2,* and *SMc03253* (17, 18). NifA is located on the symbiotic plasmid, pSymA, and is essential for nitrogen fixation. It is an enhancer-binding protein that acts in conjunction with σ^54^ to control the expression of nitrogenase (*nifH, nifDK*) and a high oxygen affinity cytochrome oxidase (*fixABCX*) (11). Two copies of FixK are also found on pSymA and are classified as *crp/fnr*-type regulators that control the expression of the denitrification genes (*nap, nir, nor, nos*) as well genes responsible for the synthesis of an oxidase with high oxygen affinity (*fixNOPQ*) (19). It is unknown why chromosomally located *proB2* and *SMc03253,* both involved in proline metabolism, are direct targets of FixJ (17, 18).

Unlike many other FixL/FixJ systems found in rhizobia, FixJ does not directly autoregulate in *S. meliloti* (Figure 8). Instead, FixK controls the expression of *fixT,* an antikinase that represses the FixLJ regulon by preventing the autophosphorylation of FixL (20). Repression by FixT is abolished in the absence of the glutamine-dependent asparagine synthetase, *asnO,* though how this occurs is still poorly understood (21). One possibility suggests that AsnO may serve a regulatory role by monitoring nitrogen balance within the cell via a metabolite, such as glutamine, and controlling nitrogen fixation accordingly (21). However, more study is required to determine how the interaction works, though it is known that neither FixT nor AsnO is required for nitrogen fixation in *S. meliloti* (22).

In addition to acting as an oxygen sensor, FixL also detects nitric oxide. Meilhoc et al. demonstrated that FixLJ along with the nitric oxide response regulator (NnrR) are involved in the *S. meliloti* response to nitric oxide exposure (23). The nitric oxide response was observed when lag phase cultures were exposed to a nitric oxide donor under aerobic conditions, as well as inside the nodule.

During symbiosis, oxygen concentrations can be as low as 5-30 nM within legume nodules, compared to the aerobic conditions in culture media where oxygen concentrations are approximately 250 μM at the start of growth (22, 24). Since most denitrification studies are performed under low oxygen tension and/or early in culture growth, we found that there is limited literature regarding denitrification in *S. meliloti* under conditions such as the higher cell population densities, which occur in the rhizosphere prior to root invasion. Previous unpublished work performed in our laboratory has revealed that a LuxR-like regulator, SMc00658, is involved in the regulation of denitrification at high cell population densities in aerobically grown cultures (Patankar AV and González JE, unpublished data). For this reason, we have changed the name of the *SMc00658* locus to the <u>a</u>erobic <u>d</u>enitrification <u>r</u>egulator, *adr*. Unlike traditional LuxR-like response regulators, Adr does not respond to the *N-*acylhomoserine lactone (AHL) molecules produced by *S. meliloti* (25). It also does not appear to be regulated by the legume symbiosis quorum sensing regulator ExpR. In the absence of Adr, we see a dramatic drop in the expression of *fixK*, *nifA*, and the downstream genes (*nap*, *nir*, *nor*, *nos*) which is remarkably similar to the expression profile of a *fixJ* mutant (17). This change in expression is independent of the presence or absence of *S. meliloti* AHLs (Patankar AV and González JE, unpublished data).

In this study, we show that the FixLJ system is actively expressed under aerobic growth conditions and that Adr is necessary for this activity. We also demonstrate that while Adr has a dramatic effect on the expression of FixK and NifA under aerobic conditions, it is not required for nitrogen fixation in the nodule.

## Results

### Adr sequence analysis

Initial sequence analysis suggests that Adr belongs to the LuxR family of transcriptional regulators. These regulators are identified by a carboxyl-terminal helix-turn-helix motif and an amino-terminal signal recognition domain. LuxR family regulators are best known for their role in quorum sensing, where they act as the signal response regulator. However, the LuxR family is large and loosely conserved, with a reported 18-25% overall sequence identity (26). *S. meliloti* has several characterized LuxR-like regulators, including the quorum sensing regulators ExpR and SinR, the motility regulators VisN and VisR, and the methyl cycle regulator NesR (27-29). Here we report a new LuxR-like transcriptional regulator in *S. meliloti,* Adr, which appears to regulate aerobic denitrification.

Though members of the LuxR family generally have very low sequence identity, 95% of known LuxR family members share nine conserved residues that are important for DNA binding and AHL signal recognition (Figure 2) (30). Sequence analysis shows that Adr only has four of these nine conserved residues, three of the DNA binding residues and only one of the signal recognition residues. Based on this analysis, it is unlikely that Adr responds to AHL signals. Additionally, when *S. meliloti* is unable to produce AHLs, there is no effect on the expression or function of Adr when compared to AHL producing cultures (Patankar AV and González JE, unpublished data).

**Figure 2.**
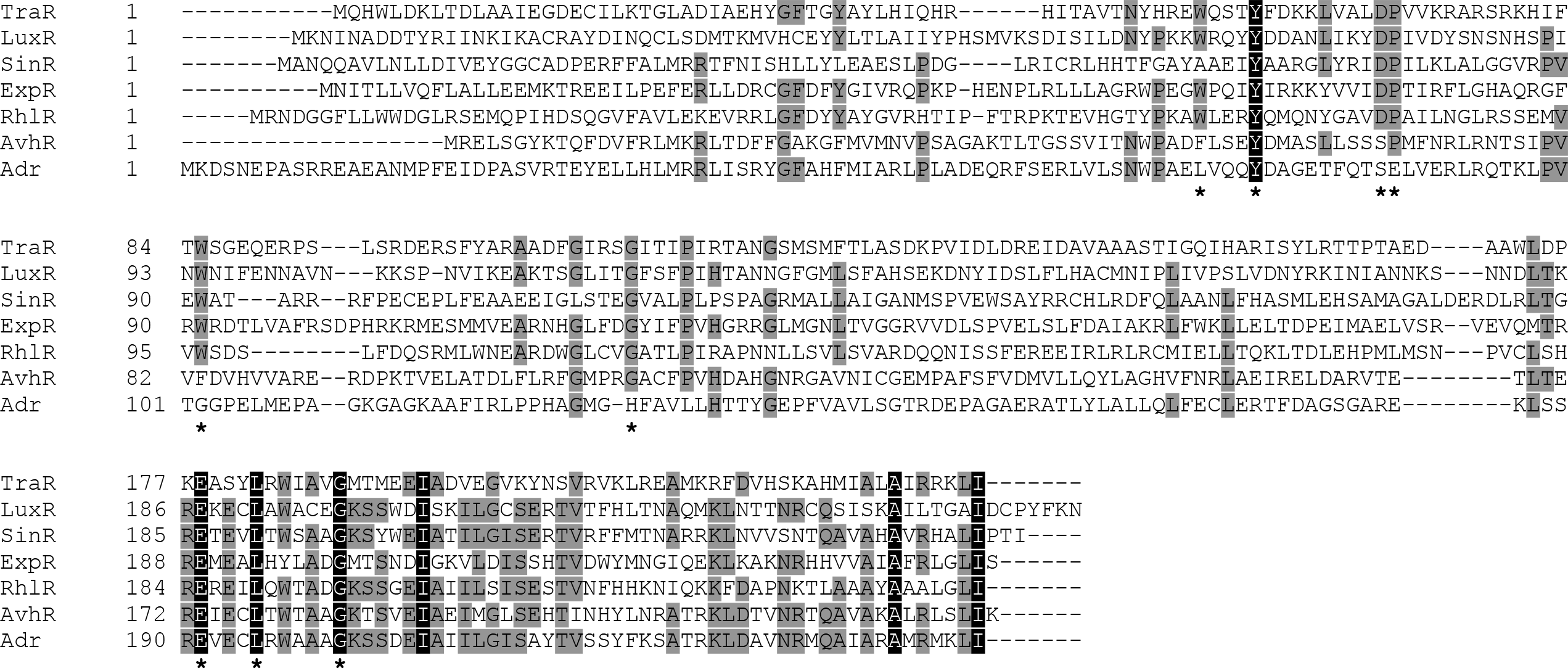
Comparison of Adr (SMc00658) to LuxR. Sequence alignment of TraR from *Agrobacterium tumefaciens,* LuxR from *Alovibrio fischeri,* SinR and ExpR from *S. meliloti,* RhlR from *Pseudomonas aeruginosa*, and AvhR from *Agrobacterium vitis* with Adr from *S. meliloti.* Grey shaded residues are highly similar between proteins and identical proteins are shaded in black. Asterisks indicate the nine highly conserved residues found in >95% of LuxR-type proteins. The alignment was performed using Invitrogen’s Vector NTI 11 software.

Among functionally characterized LuxR-like regulators reported to date, AvhR (a regulator from *Agrobacterium vitis* responsible for plant necrosis) shows the most sequence homology to Adr (28%). Consequently, this regulator also lacks the typically conserved AHL binding residues found in other classical quorum sensing LuxR type regulators (31).

### Disruption of *adr* reduces expression of denitrification and nitrogen fixation genes

To determine the regulatory role of Adr during normal growth of *S. meliloti*, we conducted a microarray analysis to compare the transcriptomic profiles of wild-type *S. meliloti* Rm8530 to Rm8530 *adr*. Since Adr is similar to quorum sensing related regulators, conditions for the microarray followed those used for other quorum sensing expression analysis microarrays (27, 32). Cultures were aerated and grown to early stationary phase (OD_600_ 1.2) before RNA was harvested. Preliminary analysis performed using the AffyMetrix GCOS software revealed that 427 genes were differentially regulated between the wild-type and the *adr* mutant (Supplemental Table S1). Of these, 247 genes were downregulated in the *adr* mutant. Within this set of downregulated genes, we were particularly interested in the genes involved in denitrification (*nap, nir, nor, nos*) and microaerobic respiration (*fix*) (Table 4). Further analysis of the microarray data revealed that the regulatory genes *fixK1, fixK2,* and *nifA* were also downregulated in the absence of Adr. Previous work on the denitrification pathway performed by Bobik et al. revealed that FixJ is the major regulator of limited oxygen response in *S. meliloti*. This response includes activating the denitrification pathway. When comparing our microarray results to those obtained by Bobik et al. we observed an 83% overlap of genes regulated by FixJ and Adr, indicating that Adr is a yet uncharacterized regulator involved in the denitrification and limited oxygen response pathways in *S. meliloti* (Table 4) (17).

**Table 1.** Strains and Plasmids.

| Strains or plasmids | Relevant characteristics | Reference or source |
| --- | --- | --- |
| *S. meliloti* |  |  |
| Rm8530 | Su47 *str-21*, *expR*<sup>+</sup>, Sm<sup>R</sup> | (57) |
| Rm8530 *adr* | *SMc00658::Tp* | This work |
| Rm8530 *fixJ* | *fixJ::Nm* | This work |
| Rm8530 *fixJ adr* | *SMc00658::Tp fixJ::Nm* | This work |
| Rm8530 *fixL* | *fixL::Nm* | This work |
| *E. coli* |  |  |
| DH5α | See source | Life Technologies |
| MT616 | MT607(pRK600) | (49) |
| S17-*λpir* | See source | (58) |
| Plasmids |  |  |
| pPCR-Script | See source, Amp<sup>R</sup> | Stratagene |
| pVIK112 | *lacZY* for transcriptional fusions, Km<sup>R</sup> | (59) |
| pJQ200SmSp | Suicide vector, *sacB*, Gm<sup>R</sup> | (25) |
| pMB419 | Vector carrying Hygromycin cassette | (60) |
| p658 | pPCR-Script carrying *SMc00658* | This work |
| p658Tp | pPCR-Script carrying *SMc00658* disrupted with Tp | This work |
| pJQ658Tp | pJQ200 SmSp carrying *SMc00658::Tp* | This work |
| pVIKfixK1 | pVIK112 carrying *fixK1* fragment | This work |
| pK19fixK2 | pK19mobΩHMB carrying *fixK2* fragment | Anke Becker |
| pK19fixK2Hy | pK19mobΩHMB carrying *fixK2::Hy* | This work |

**Table 2.**
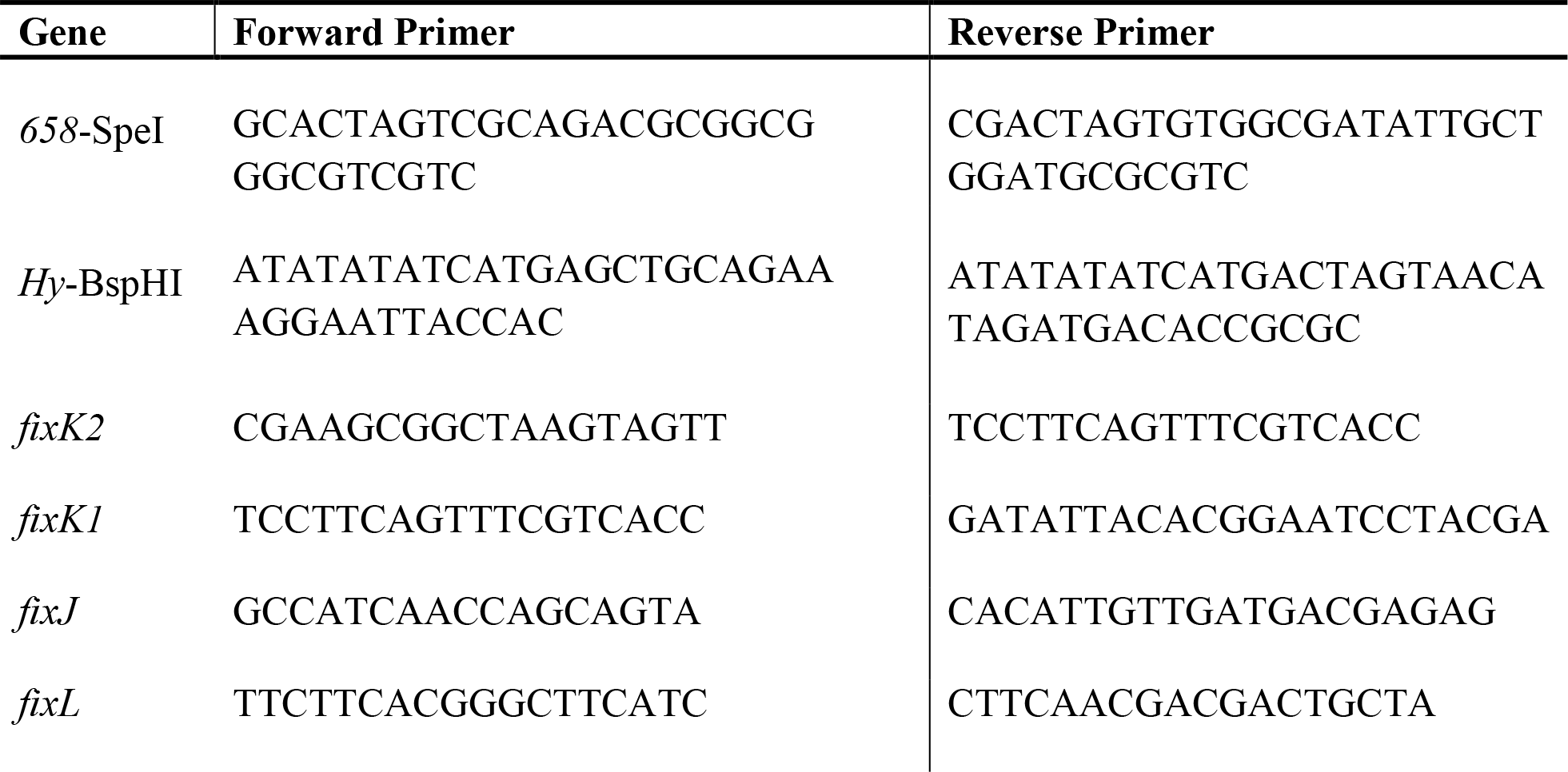
Primers used for mutant construction.

**Table 3.**
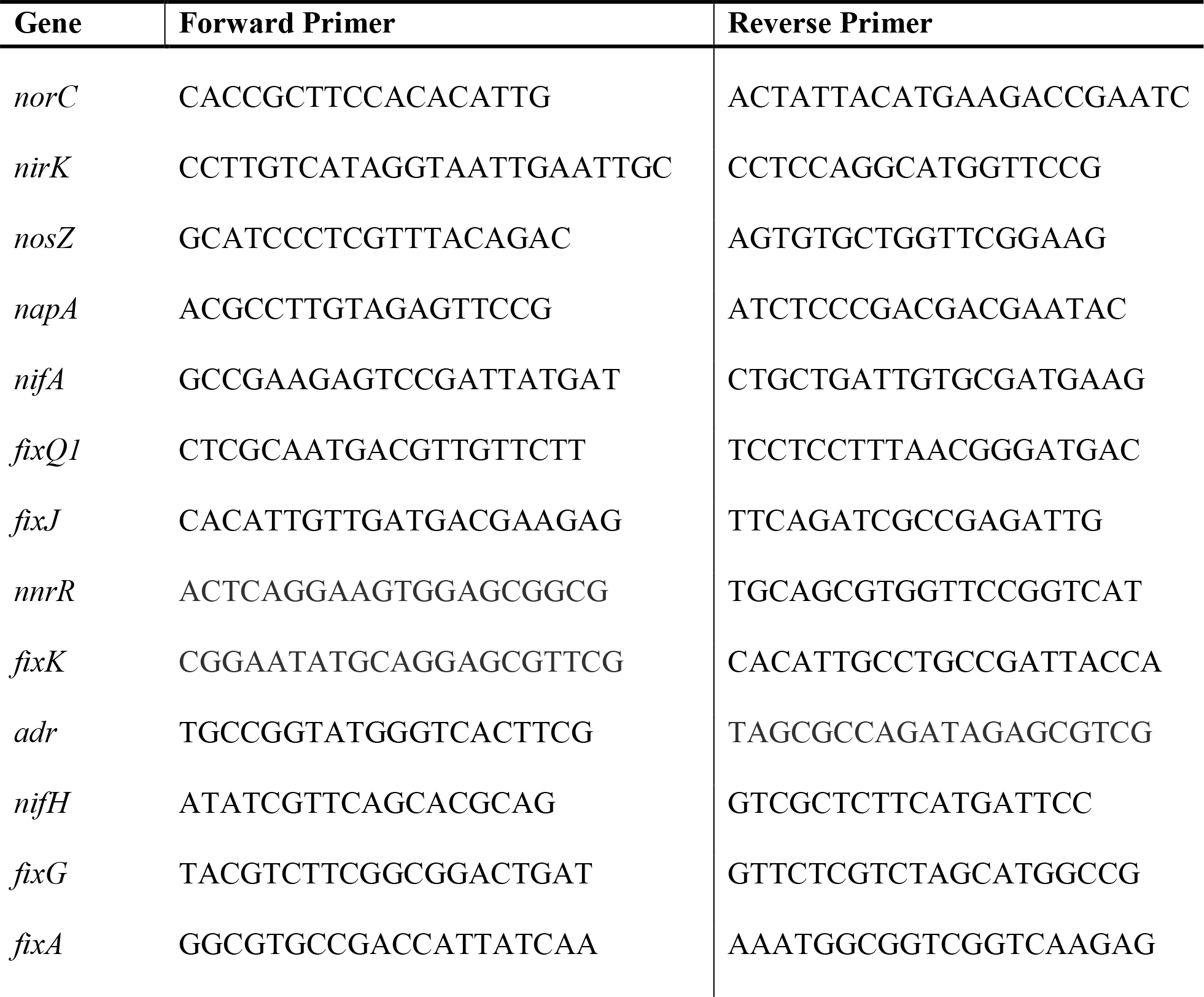
qRT-PCR Primers.

**Table 4.**
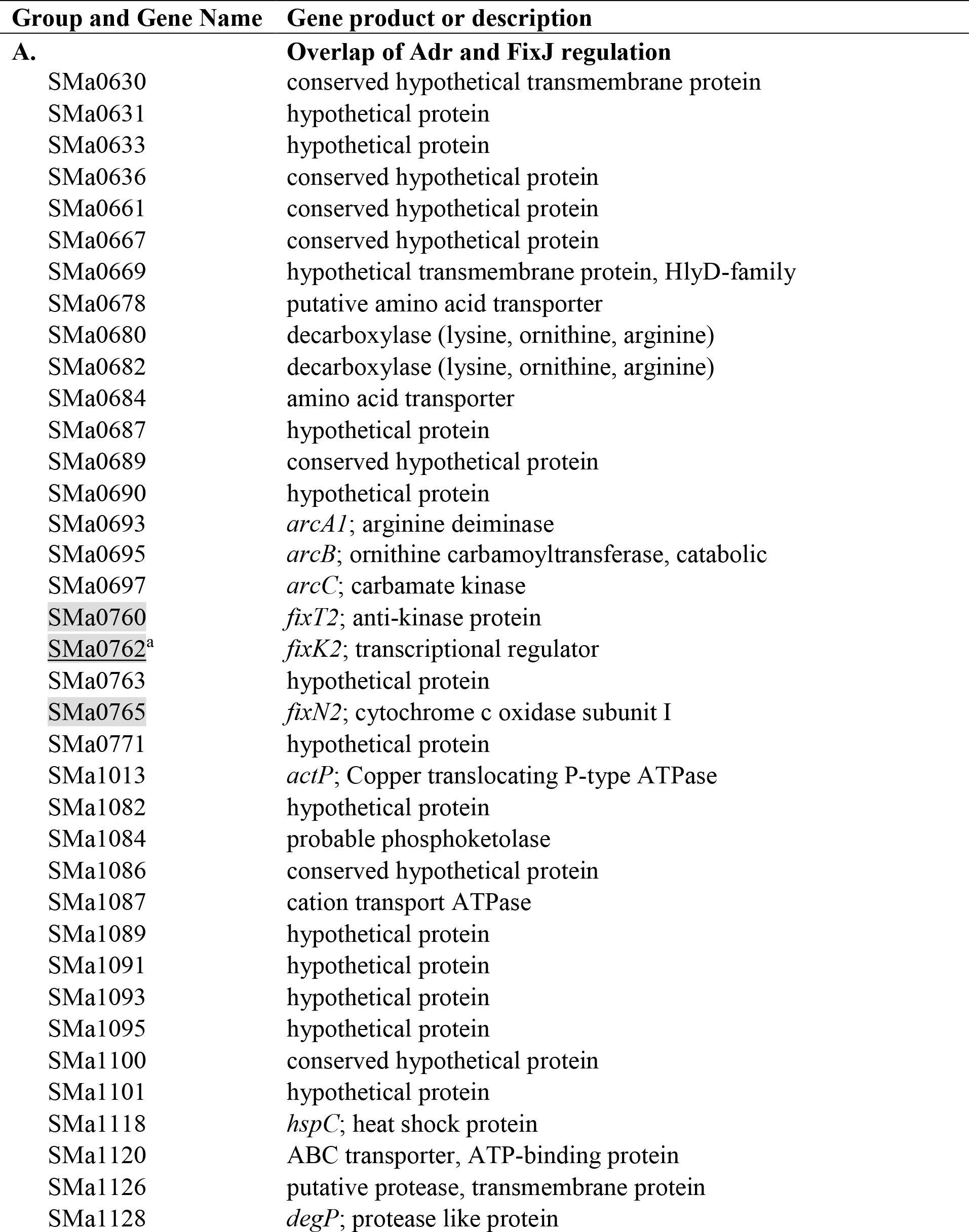
Comparison of the FixJ by Bobik, et al. and the Adr microarray.

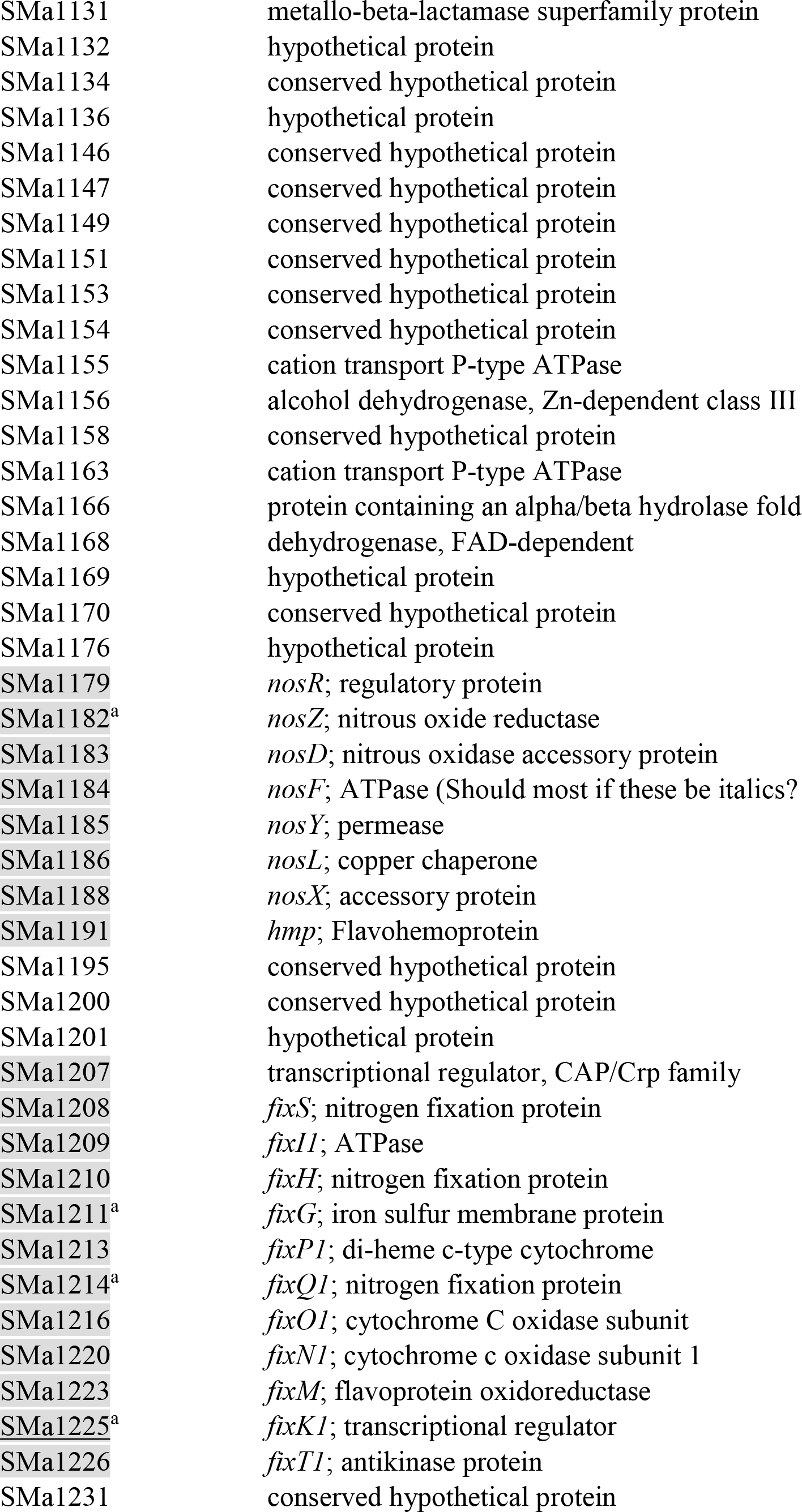

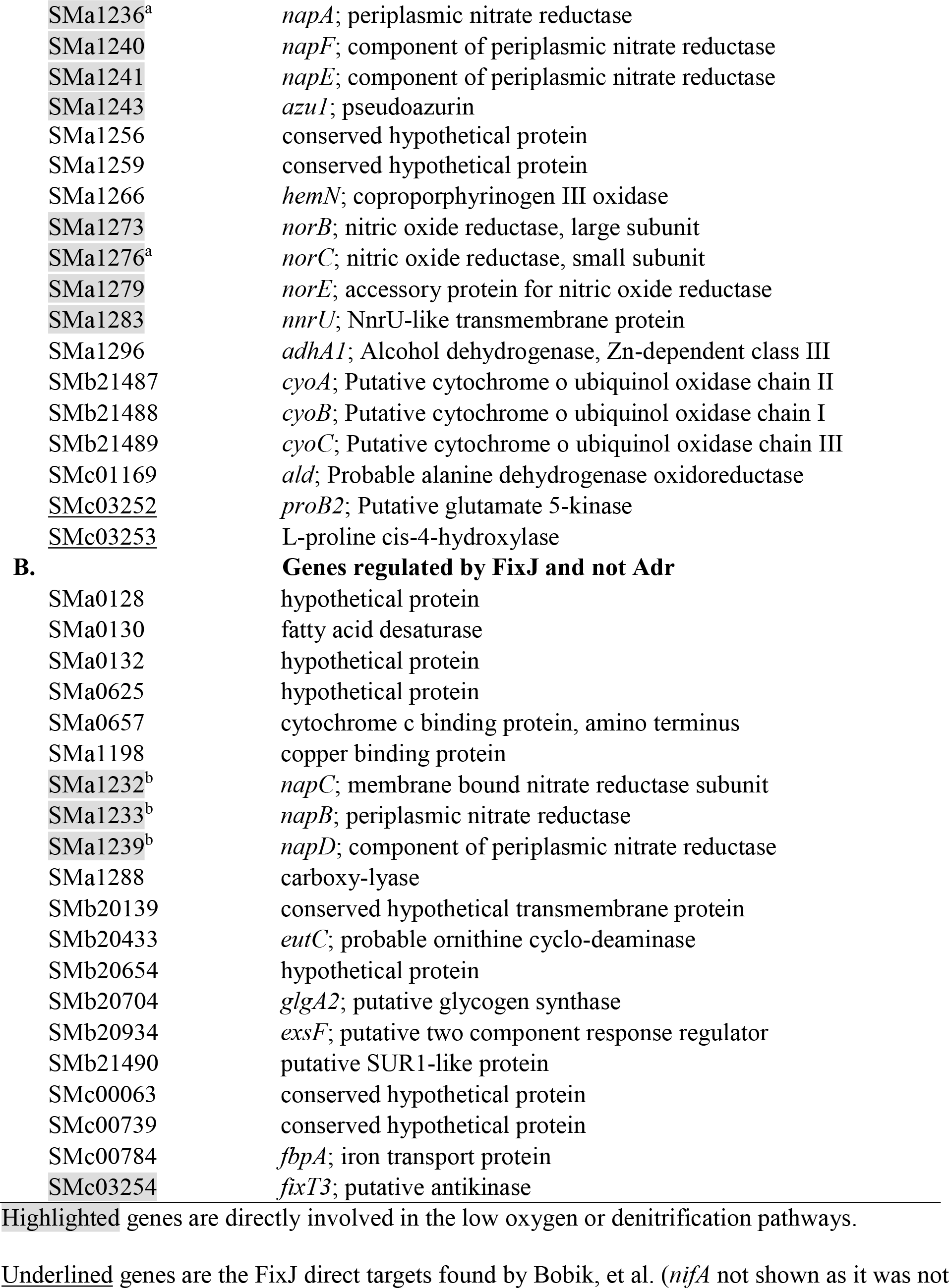

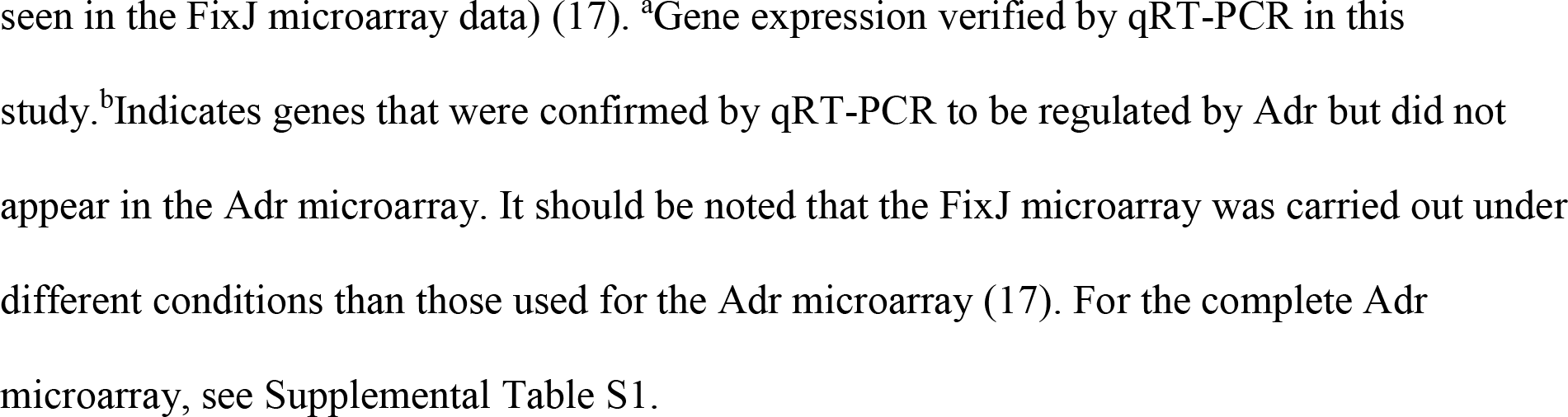

To verify our microarray results, several genes involved in the denitrification pathway were selected for quantitative real time-PCR analysis (Figure 3). Results are represented as fold change, which is 2^ΔCt^, where ΔCt equals the difference in expression of the wild-type from the mutant. *napA, nirK, norC,* and *nosZ* are structural subunits of nitrate reductase, nitrite reductase, nitric oxide reductase, and nitrous oxide reductase, respectively. Genes were selected based on M-value (>1.5). The results of this analysis indicate a significant downregulation (5- to 300-fold change) in each step of the denitrification pathway in the absence of Adr (Figure 3a). We also included *fixK* and *nifA* in this analysis since these two regulators also appeared in our microarray data. The reduction of *fixK* (16-fold) and *nifA* (42-fold) expression seen in Figure 3a in the *adr* mutant is typically seen in a *fixJ* mutant, indicating that Adr impacts these genes even in the presence of FixJ. The decrease of FixK is likely causing the downstream effect in the denitrification genes.

**Figure 3.**
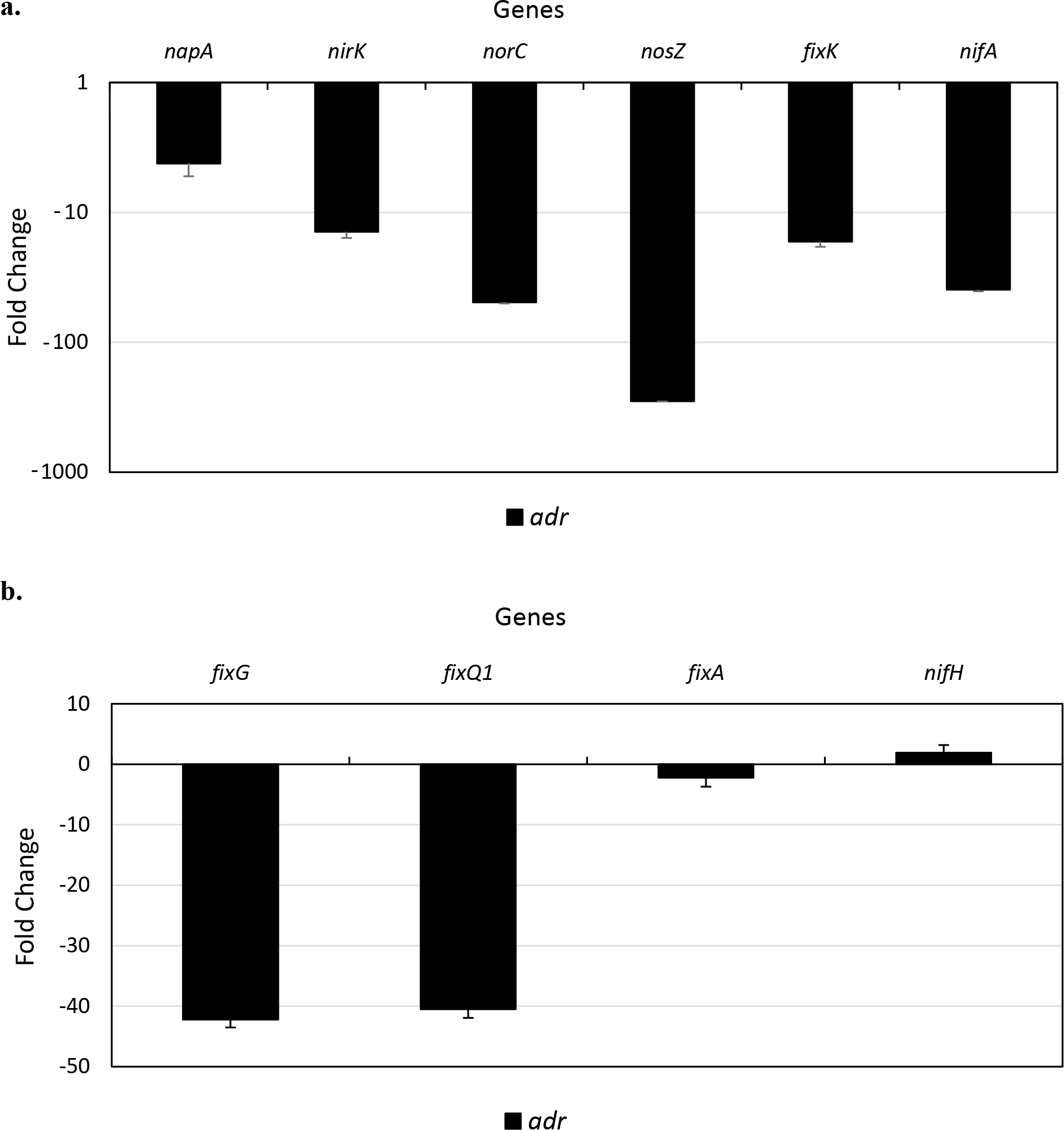
Relative expression of the *adr* mutant compared to wild-type strain Rm8530. Expression was measured using qRT-PCR and relative transcript levels are displayed as fold change between the wild-type Rm8530 and the *adr* mutant. Negative values indicate downregulation of the denoted gene in the mutant strain in comparison to the wild-type, while positive values represent the inverse. Results are the average of three independent biological replicates, error bars are present and represent the standard deviation between three samples. *SMc00128* was used as an internal control (52).

We next measured the effect of Adr on other FixK dependent genes, including the two subunits of high oxygen affinity cytochrome oxidases, *fixG* and *fixQ1* (Figure 3b) (33). As was the case with the denitrification pathway genes, the lack of Adr lead to a decrease in *fixG* and *fixQ1* expression (39-fold and 42-fold respectively), confirming that FixK is downstream of Adr when regulating genes related to the denitrification and limited oxygen response pathways.

We also measured *nifA* expression since, like FixK, it is controlled by FixJ. As was the case with *fixK,* the removal of Adr leads to a decrease in expression of *nifA* (Figure 3a). However, unlike with FixK, we found that genes downstream of NifA, such as *nifH* or *fixA,* were not affected by the presence or absence of Adr (Figure 3b). This is unsurprising since previous studies have shown NifA to be sensitive to oxygen levels higher than those found in the nodule; though the *nifA* gene is being expressed, its translated form is not active under our aerobic growth conditions (3).

Though not seen in our microarray data, we also tested the effect of Adr on *nnrR*, the nitric oxide response regulator. This regulator induces the transcription of nitrite reductase (*nirKV*) and nitric oxide reductase (*norECBQD*) in the presence of nitric oxide. While FixLJ also responds to nitric oxide, NnrR is not in the FixJ regulon (23). Under our conditions, we saw no differential expression of *nnrR* (data not shown).

Since the expression of both *nifA* and *fixK* are impacted by the absence of Adr, it appears that Adr is acting in a manner that is either parallel and/or in conjunction with FixJ.

### Adr and FixJ regulate denitrification in tandem

Previous studies of *S. meliloti* denitrification have revealed FixJ as the major denitrification regulator (Figure 8) (17, 34). These studies approached denitrification regulation by examining expression under microoxic (<2% oxygen) conditions, comparable to those found before the cells enter the nodule as well as after the cells differentiate into nitrogen fixing bacteroids. In our study, we show that many of the same genes regulated by FixJ also show differential expression in the absence of Adr in aerobically grown cells. Though not a direct comparison due to the difference in conditions, we found that 83% of genes reported by Bobik, et al. to be regulated by FixLJ are also differentially expressed in the Adr mutant (17). Additionally, it was also noted that the five genes directly regulated by FixJ (*fixK1*, *fixK2*, *nifA*, *proB2*, and *SMc03253*) were also found to be regulated by Adr (Table 4) (18).

In light of these findings, we generated a *fixJ* mutant to determine if the FixLJ system is active under the same conditions as Adr. qRT-PCR analysis of the *fixJ* mutant revealed that the FixJ is actively influencing denitrification expression in aerated stationary phase cultures (Figure 4a). Direct targets of FixJ such as *fixK* and *nifA* were downregulated (approximately 1000-fold) in the *fixJ* mutant; downregulation was also see in the indirect targets including the denitrification (100- to 1000-fold) and microoxic respiration genes (65- to 70-fold) (Figure 4b and 4c). However, as with the *adr* mutant, we did not see a large change in expression of genes controlled by NifA (a direct target of FixJ) such as the nitrogenase structural gene *nifH* or a subunit of another oxidase, *fixA* (Figure 4c). This supports data from other laboratories that found that nitrogenase and its auxiliary proteins are only expressed in bacteroids (3).

**Figure 4.**
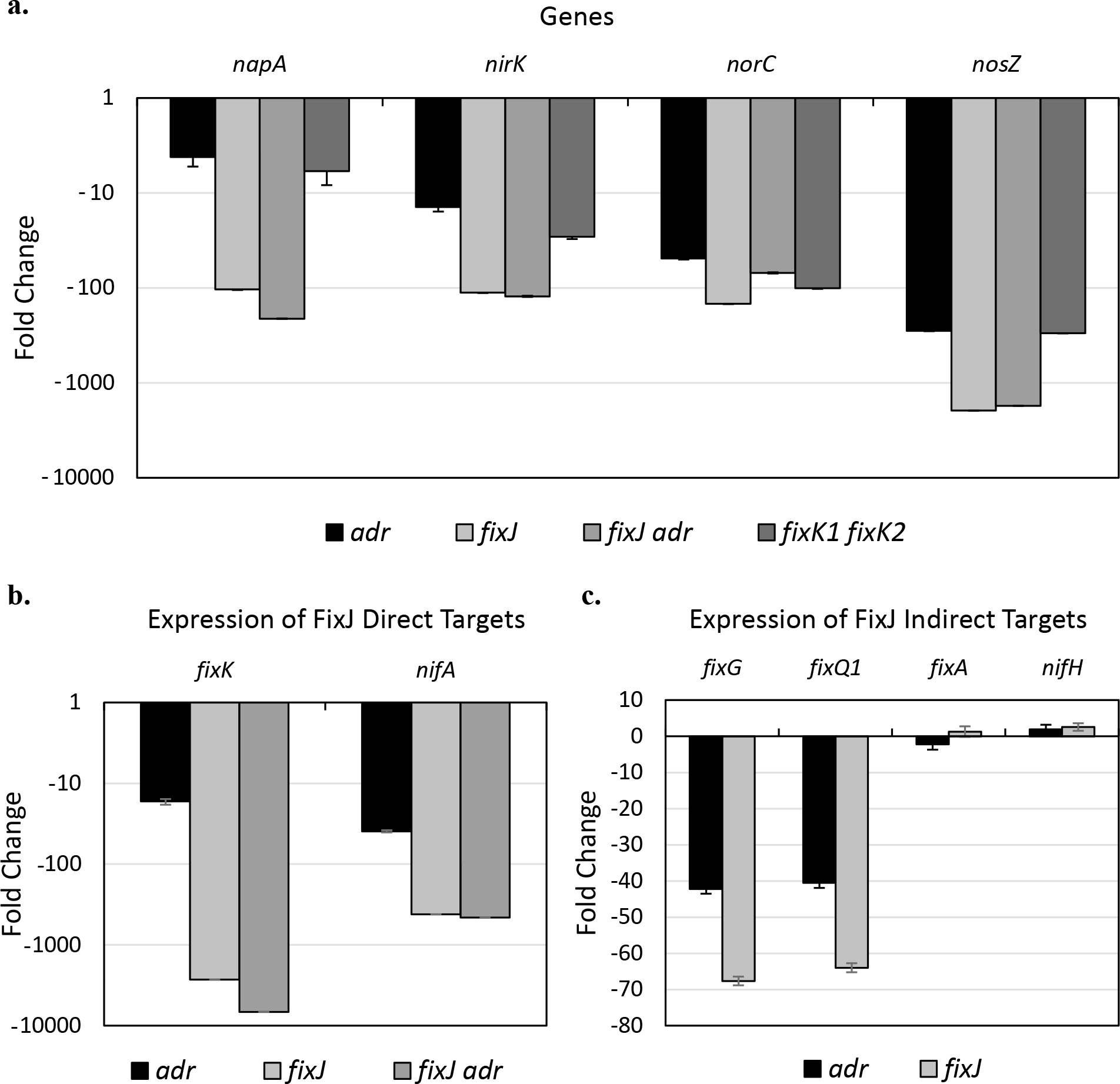
Aerobic expression of denitrification and other FixJ regulated genes. **a)** The expression of denitrification genes from four mutant strains (*adr, fixJ, fixJ adr*, and *fixK1 fixK2*) compared to wild-type Rm8530 expression levels. **b)** Direct targets of FixJ, *fixK* (total expression of both genes) and *nifA*. **c)** Comparison of expression between four indirect FixJ targets. *adr* mutant data is the same as that seen in Figure 3, repeated for clarity. Results are the average of three independent biological replicates. Error bars are present and represent standard deviation. *SMc00128* was used as an internal control (52).

The presence of an active FixJ revealed the possibility that *adr* is either regulated by the FixLJ system or regulated independently but affecting the same genes. To this end, we measured the expression of *adr* in the absence and presence of *fixJ*. We also tested the reverse to determine if *fixJ* expression depends on the presence of *adr.* In both cases we saw no differential expression between the wild-type and mutant backgrounds (data not shown).

A *fixJ adr* double mutant was also constructed to determine if there is an additive effect on expression by Adr and FixJ. Expression levels of *fixK, napA, nirK, norC,* and *nosZ* were compared between the *adr*, *fixJ*, and *fixJ adr* double mutant. Results show that there is no compounding effect by introducing an *adr* mutation into a *fixJ* mutant (Figure 4a).

To verify that Adr is working at the FixJ level, we measured the expression of the denitrification genes in a *fixK* double mutant. The results confirm that when FixK is not expressed, which we suspect occurs in an *adr* mutant, the denitrification genes are not expressed (Figure 4a).

### Expression of *adr* during growth

Past studies have focused on the activity of FixJ and as a consequence experiments were performed under conditions that lead to FixLJ activation. Therefore, previous work was generally done early in the growth cycle (OD_600_ 0.2-0.5) and under microoxic (2% O2) conditions. To determine if growth phase had an effect on Adr expression, we performed additional expression measurements during lag (OD_600_ 0.2) and mid exponential (OD_600_ 0.8) growth which was compared to our stationary phase data (OD_600_ 1.2). Results show that there is very little change in expression of the *napA*, *nirK*, *norC*, and *nosZ* genes during lag growth when wild-type was compared to the *adr* mutant. However, activation of denitrification gene expression occurs during mid exponential phase and continues into stationary phase (Figure 5). We also analyzed the expression of *adr* over time and found that there is no difference in expression of the gene in wild-type Rm8530 between the three time points tested (Table 5).

**Table 5.** Relative transcript levels of *adr* over time, represented as Ct value*.

| **Strain** | **Lag**<br>**(OD<sub>600</sub> 0.2)** | **Mid-exponential**<br>**(OD<sub>600</sub> 0.8)** | **Early Stationary**<br>**(OD<sub>600</sub> 1.2)** |
| --- | --- | --- | --- |
| **Rm8530** | 34.74 ± 0.48 | 33.34 ± 0.46 | 34.64 ± 0.34 |

\*Ct value represents the arbitrary cycle where PCR product is detected. Values closer to 40
indicate lower amounts of starting transcript.

**Figure 5.**
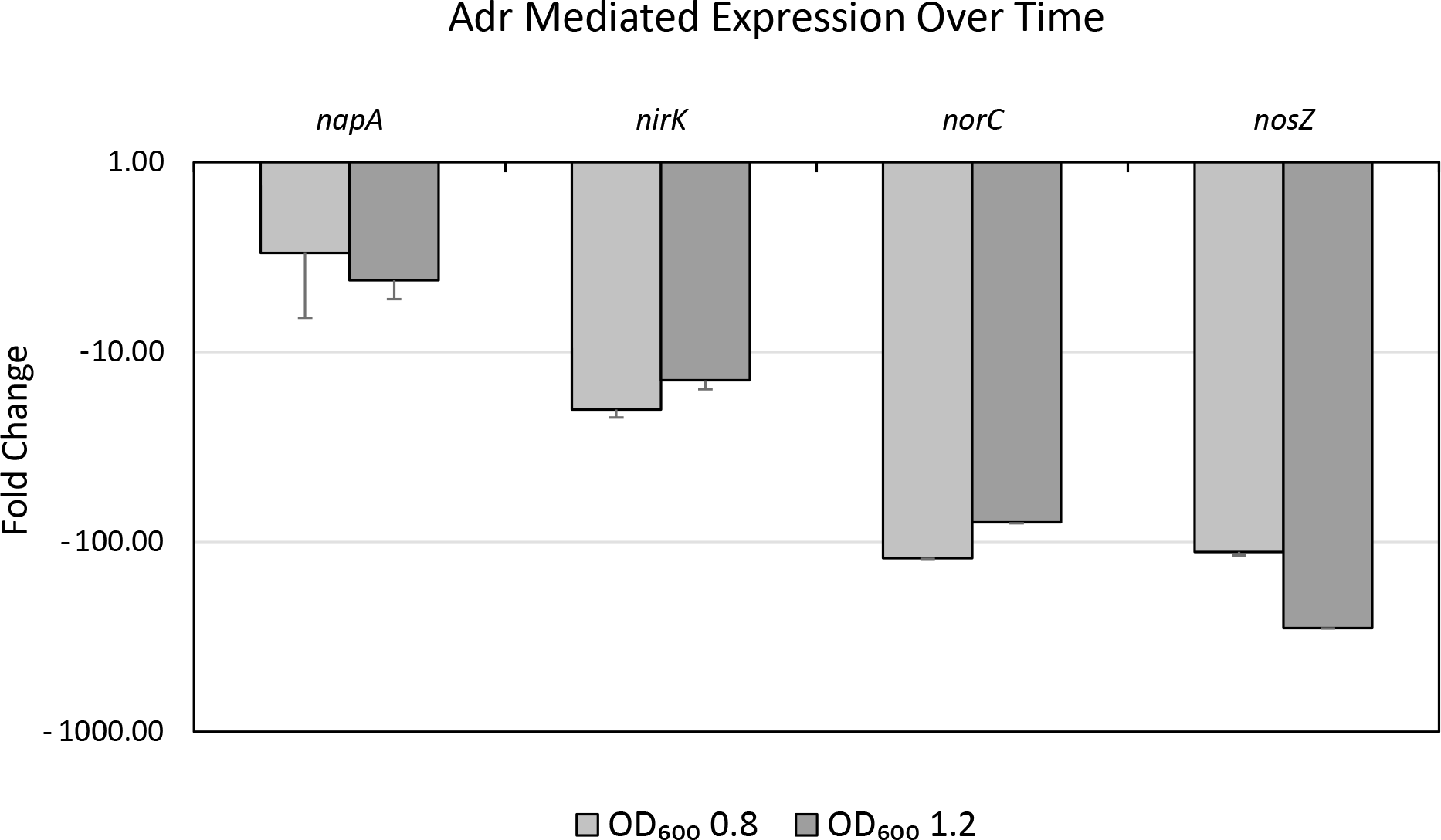
Expression of denitrification genes in an *adr* mutant over time. Cells were grown to the indicated optical densities (measured at 600 nm). Expression was measured in three independent biological replicates per density and all are compared to the wild-type expression value. Error bars represent standard deviation. No difference in expression was observed at OD_600_ 0.2. *SMc00128* was used as an internal control to normalize values (52).

Since differential expression due to Adr was detected during exponential growth, we next attempted to determine if a media soluble signal was responsible for this change. Spent media was harvested from early stationary phase cultures and used to grow fresh cultures of Rm8530 and Rm8530 *adr* to OD_600_ 0.2. While no difference in expression was seen between cultures using fresh versus spent media, we do not rule out the possibility of a signal produced by the cell involved in the activation of Adr (data not shown).

### Effect of the absence of FixL

As discussed previously, FixL is the oxygen sensing component of the FixLJ system. Our data presented here shows that there is a relationship between Adr and FixJ under aerobic conditions. To determine if FixL has a role in this relationship by interacting with Adr, a *fixL* mutant was generated and expression of *adr* was measured. Expression of *adr* was not affected by the absence of FixL in stationary phase (data not shown).

### The Adr mutant shows decreased survival under oxygen limitation in the presence of nitrite

Since Adr enhances expression of the denitrification pathway under aerobic conditions, we measured the effect of Adr during aerobic growth. Rm8530 (wild-type) and Rm8530 *adr* were grown in TYC media supplemented with either 10 mM KNO_3_, or 5 mM NaNO_2_ was. The growth rate was similar between all strains under all three conditions (Figure 6a). Viable counts were performed on aerobically grown cultures after 72 hours to determine survival of *S. meliloti* under conditions listed above. No difference in survival was observed (Figure 6b).

**Figure 6.**
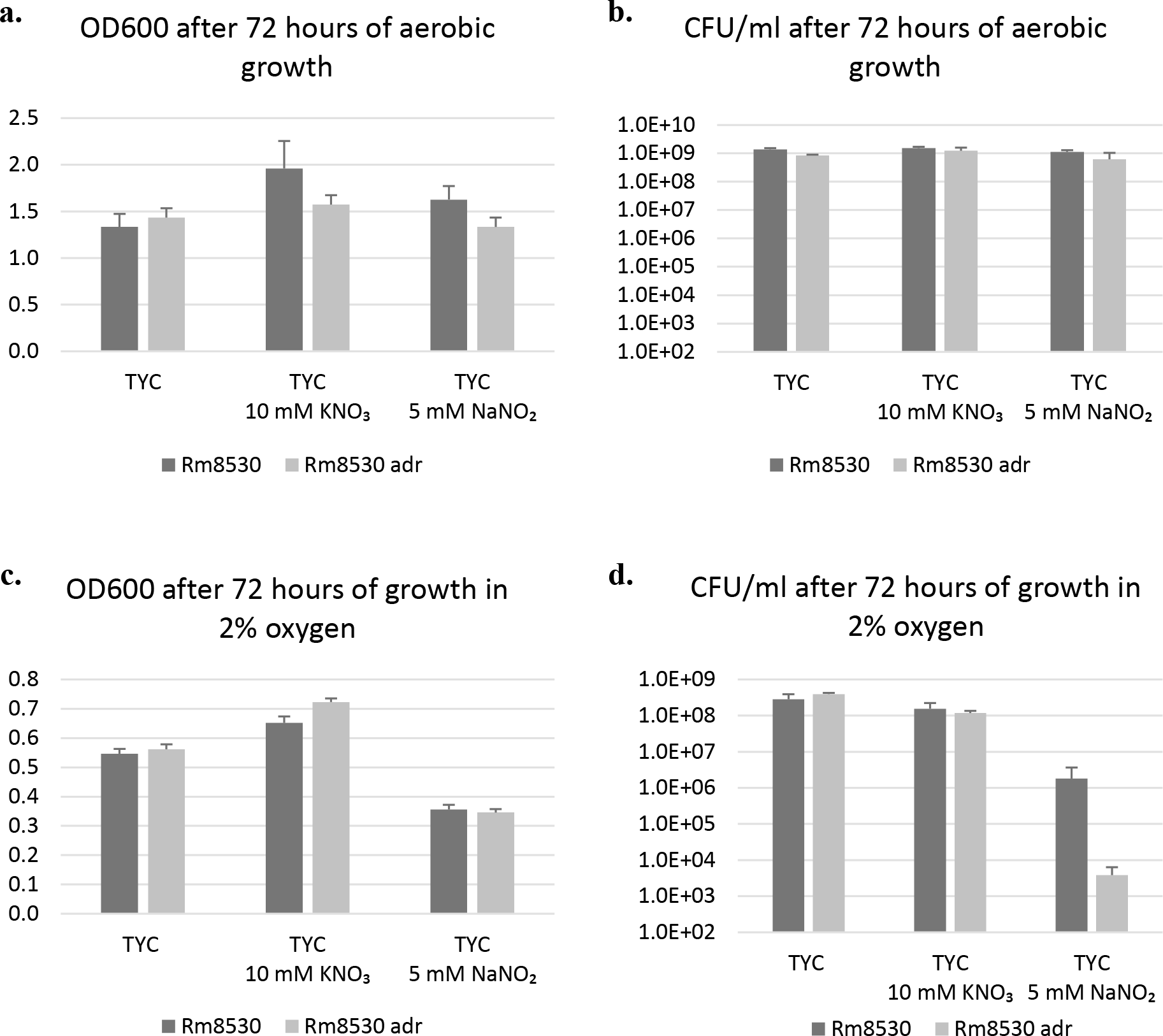
Growth and survival of the Adr mutant. Growth measurements of wild type Rm8530 and the mutant Rm8530 *adr* were conducted in triplicate under aerobic and microaerobic conditions as described previously

Since no difference in growth or survival was observed during aerobic growth conditions, we also conducted the same growth and survival assay under microaerobic conditions. Cultures were grown in LB/MC for two days then diluted to OD_600_ 0.2 in TYC before microaerobiosis was induced by filling the headspace of the vial with a mixture of 2% oxygen and 98% argon gas. Figure 6c shows that, while growth appeared to be similar between the wild-type and mutant under all three conditions, the number of viable colony forming units dropped significantly when the *adr* mutant was grown in media containing nitrite (Figure 6d). Anaerobic conditions were also tested but no growth or denitrification was seen after seven days of incubation (data not shown).

### The nitrate reductase activity in *S. meliloti*

The *S. meliloti* nitrate reductase is encoded by the *nap* genes, which are controlled by FixJ, though expression of these genes in *S. meliloti* was thought to be constitutive regardless of oxygen concentration (2, 35). The microarray data and qRT-PCR analysis presented here indicate otherwise. We have shown that the *nap* genes are expressed by wild-type *S. meliloti* during aerobic growth, but that the expression is controlled by both FixJ (through FixK) and Adr (Figure 4a). Previous studies have demonstrated that under microaerobic conditions, the *S. meliloti* strain Rm2011, a strain very similar to Rm8530, has an active nitrate reductase and can convert nitrate to nitrite during growth (36). To determine if the transcripts of the *nap* genes are being translated into a functional enzyme in our strains, we performed assays using methyl viologen to determine the Rm8530 and Rm8530 *adr* nitrate reductase activities. Cells grown aerobically or microaerobically in nitrate free minimal media were tested. In both the aerobically grown wild-type and *adr* mutant samples, no change in absorbance was observed after adding 10 mM of KNO_3_, indicating that the conditions tested, these strains do not produce detectable levels of nitrate reductase. The same results were seen in cells grown in 2% oxygen (data not shown). However, when the cells were grown for 72 hours such as in Figure 6, low amounts of nitrite were detected qualitatively (via nitrite detection strips) in cultures of both Rm8530 and the *adr* mutant grown in 10 mM KNO_3_, indicating that there is a low level of nitrate reductase activity.

### The Adr mutant has reduced nitrite reductase activity

*S. meliloti* encodes a copper-containing nitrite reductase, *nirK*, whose enzyme product is found in the periplasm (35, 37). As it appears that *S. meliloti* is incapable of utilizing nitrate for the first step of denitrification, we next tested whether nitrite could be used to initiate the reaction. Under microaerobic conditions, wild-type Rm8530 showed a growth and survival deficiency (Figure 6c and 6d). However, this growth defect was not as dramatic as that seen in the *adr* mutant. This led us to believe that Rm8530 is capable of using nitrite to initiate denitrification, and that the inability to do so prevents the mutant from removing this toxic compound from the environment, resulting in slower growth and poor survival. To test the nitrite reductase activity, we performed the same methyl viologen assay as described above, using 10 mM sodium nitrite as the substrate. We found the nitrite reductase activity of wild-type cells to be ten times higher than that of the *adr* mutant when the cells were grown aerobically (Table 6). This observation matches our qRT-PCR data in that the expression of *nirK* is reduced in the Adr mutant when grown under aerobic conditions, but not as reduced as the expression levels of *nirK* in the FixJ mutant (Figure 4).

**Table 6.** Specific activity of nitrite reductase estimated by methyl-viologen reduction.

| Strain | Nitrite reductase activity<br>Aerobic | Nitrite reductase activity<br>Microaerobic |
| --- | --- | --- |
|  | μmol/min/mg of protein | μmol/min/mg of protein |
| **Rm8530** | 0.372 ± 0.031 | 0.935 ± 0.076 |
| **Rm8530 *adr*** | 0.042 ± 0.004 | 0.442 ± 0.049 |

Microaerobically grown wild-type cells also exhibited higher nitrite reductase activity than the *adr* mutant, though the mutant was still capable of reducing nitrite (Table 7). We also observed that under microoxic conditions, cells lacking *adr* can express nitrite reductase at a level comparable to wild-type under aerobic conditions (Table 6). We predict that this change in expression of nitrite reductase from aerobic growth to microaerobic growth is due to the activation of the FixLJ system under microoxic conditions. However, it appears that for optimal denitrification under these conditions using nitrite as the substrate, both FixJ and Adr are necessary.

### Absence of Adr results in a decrease in competitiveness during *Medicago sativa* symbiosis

Successful symbiosis and nitrogen fixation requires *S. meliloti* to not only perform the appropriate symbiotic functions at the right time, but it must also compete against other organisms in the rhizosphere for resources and hosts. To determine the symbiotic and competitive characteristics of an *adr* mutant, plant nodulation assays were performed. *Medicago sativa* inoculated with either Rm8530 or Rm8530 *adr* were equally proficient at forming nitrogen fixing nodules (Figure 7a). Since it has been shown that FixJ is a regulator of symbiosis, we also included a *fixJ* mutant and the *fixJ adr* double mutant in the nodulation assay to ensure that any effect that Adr has on nodulation was not obscured by an active FixJ. However, as shown in Figure 7a, there is no significant difference between plants inoculated with Rm8530 *fixJ* and those inoculated with the double mutant. Both sets of plants exhibited the classic Fix-phenotype of poor growth and a severe decrease in the number of nitrogen fixing nodules, which is typical of FixJ mutants.

**Figure 7.**
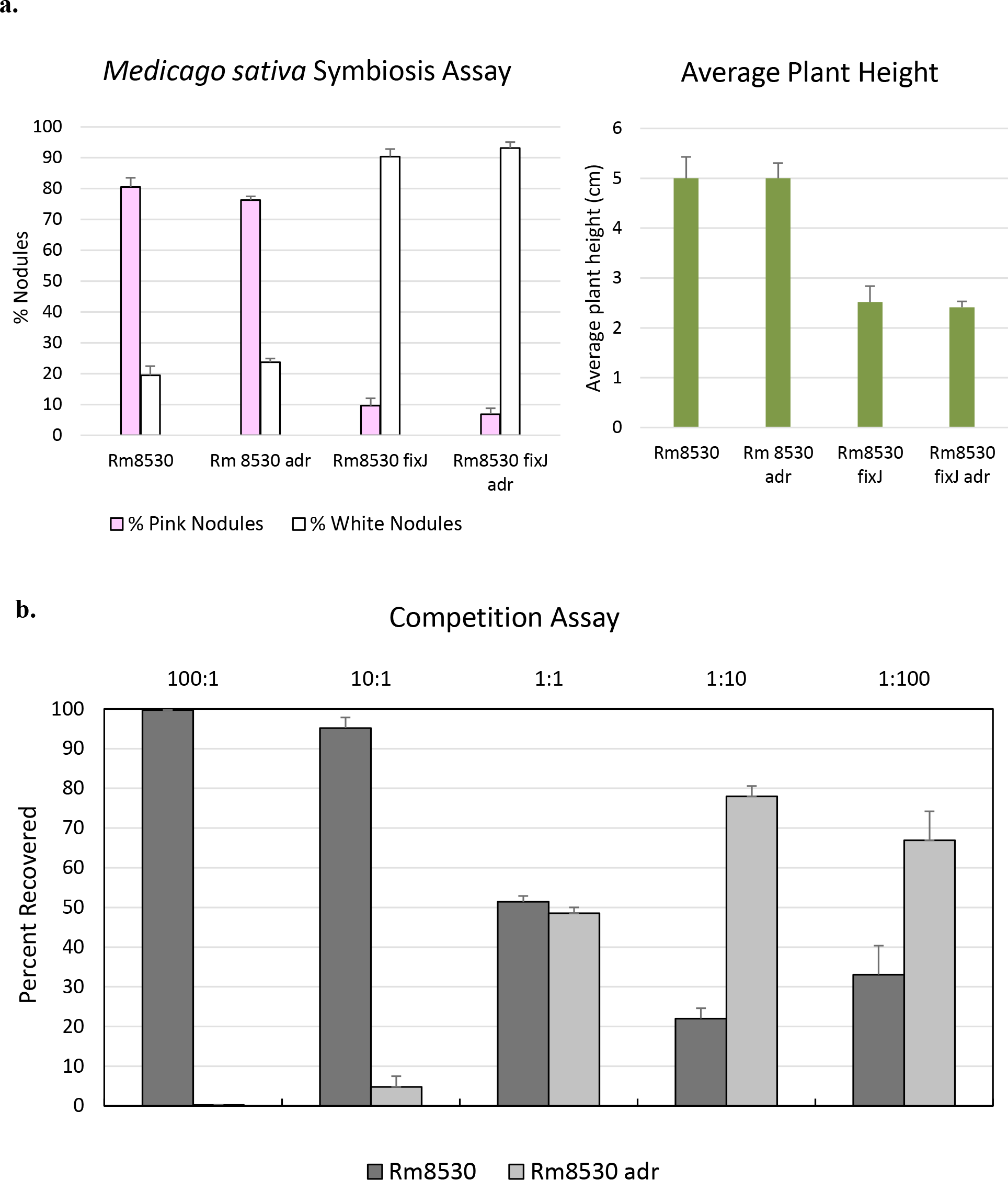
Symbiosis and competition assays. The *adr* mutant was capable of establishing symbiosis at levels comparable to the wild-type Rm8530, but was unable to compete with wild-type for nodule occupancy when both strains were co-inoculated. **a)** *Medicago sativa* cv. Iroquois inoculated with wild-type and the *adr* mutant were proficient at forming nitrogen fixing nodules and the plants were healthy. Strains deficient in *fixJ* were unable to establish nitrogen fixing nodules and resulted in plants that were not as healthy as the wild-type and *adr* mutant plants. The results are the average of three independent experiments and the standard deviations are shown. **b)** Plants were co-inoculated with varying proportions of Rm8530 and the *adr* mutant, represented on the X-axis. The percentage of bacteria recovered is shown on the Y-axis.

**Figure 8.**
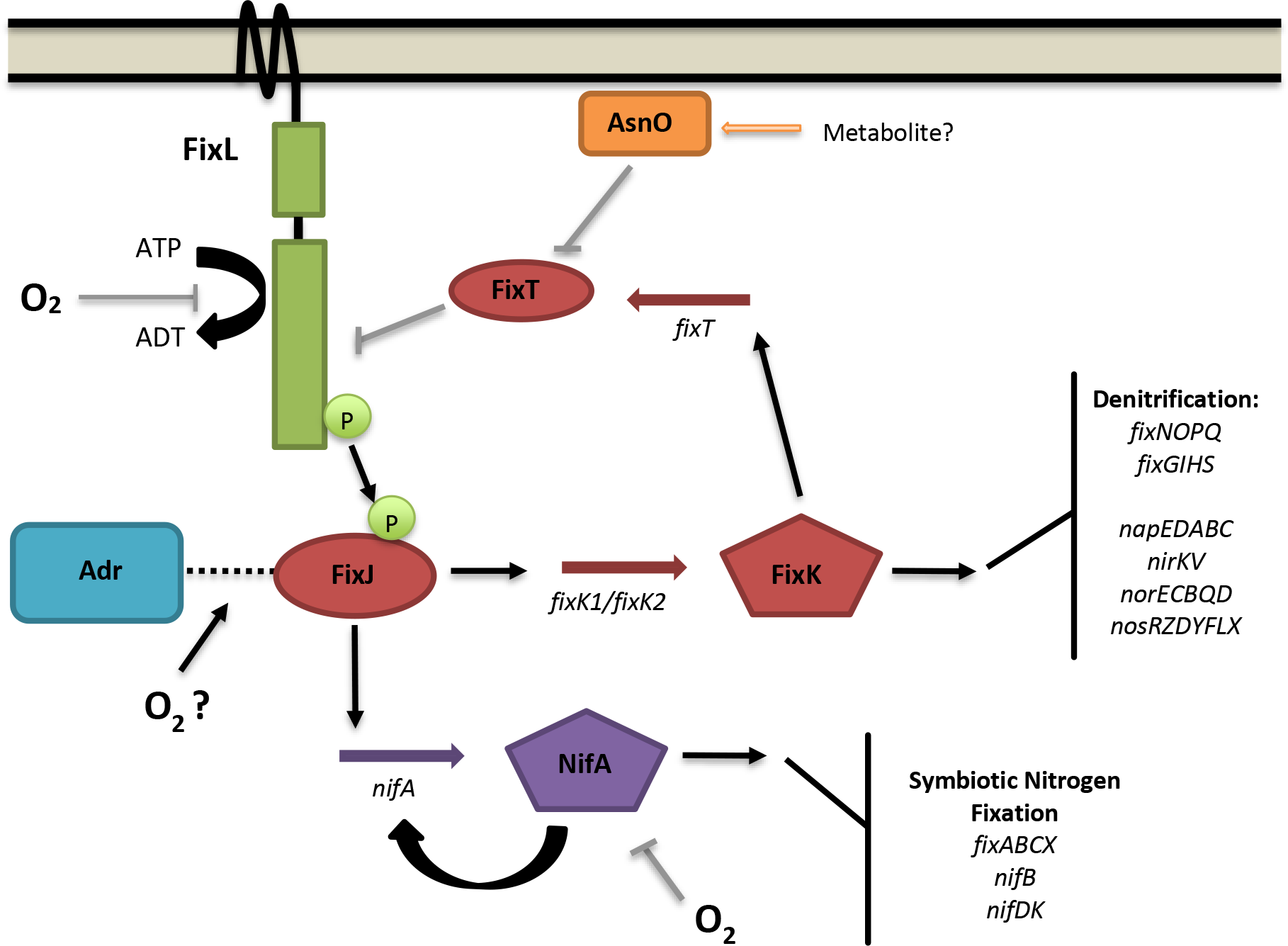
Model for regulation of denitrification and nitrogen fixation *S. meliloti*. Revised comprehensive model for the regulation of denitrification and nitrogen fixation in the presence and absence of oxygen. Grey lines indicate an inhibitory effect while black arrows indicate a positive effect. In the presence of oxygen, the autophosphorylation of FixL is inhibited, leading to a low level of phosphorylated FixJ in the cell. Adr interacts with FixJ (dotted line) and promotes the expression of the denitrification regulator FixK and the nitrogen fixation regulator NifA. FixK goes on to induce expression of the denitrification genes, as well as the antikinase FixT, which further inhibits the autophosphorylation of FixL. NifA is oxygen sensitive and is deactivated, therefore preventing the expression of the symbiotic fixation genes. When the oxygen concentration is low, FixL can autophosphorylate and transfer the phosphate group to FixJ. In the phosphorylated form, FixJ can activate the expression of the denitrification and nitrogen fixation genes without the help of Adr. AsnO acts to inhibit the antikinase activity of FixT, allowing this feedback loop to continue until environmental conditions change. Figure adapted from Terpolilli, et al. (16).

While there appears to be no difference between wild-type Rm8530 and Rm8530 *adr* invasion efficiency in monoculture, we also assessed the ability of these strains to compete for invasion when co-inoculated. Wild-type Rm8530 and the *adr* mutant were mixed in a range of ratios (100:1, 10:1, 1:1, 1:10, and 1:100) and applied to plants. After harvesting bacteria from the root nodules, we found that that Rm8530 was able to out-compete the *adr* mutant during nodulation (Figure 7b). This indicates that while Adr is not essential for nitrogen fixing nodules to form, it is beneficial to *S. meliloti* during growth in the rhizosphere.

## Discussion

Understanding the process of denitrification and its regulation is an ongoing task. While the machinery of denitrification and nitrogen fixation have been known for some time, determining how these pieces fit together under a complex regulatory web has proved to be a challenging undertaking. Though most denitrification regulatory elements are very similar, such as a sensory two-component system and *crp/fnr*-type regulators, interactions between these components vary between denitrifiers (2). Members of the same regulatory family perform different roles in various genetic backgrounds and target different respiratory systems while regulating denitrification (2). For example, in *S. meliloti* and *Bradyrhizobium japonicum,* FixL senses low oxygen tension and activates a regulatory cascade through FixJ that leads to the expression of the denitrification and nitrogen fixation pathways (38). Additionally, both *S. meliloti* and *B. japonicum* possess duplicated copies of the *fixK* gene, a *crp/fnr*-type regulator. However, in *S. meliloti* both copies of *fixK* activate the same set of promoters, while in *B. japonicum* FixK2 is essential for denitrification and nitrogen fixation, as well as regulating the expression of FixK1 (16). In *Pseudomonas* species, a variant of FixLJ, the NarXL system, controls denitrification and NarL acts as the transcriptional activator of the nitrate reductase genes (39, 40).

Unlike the typical model denitrifiers, genes for the denitrification pathway in *S. meliloti* are located on the symbiotic plasmid pSymA instead of the chromosome (2, 33). Generally, rhizobia are considered to have acquired nitrogen fixing capabilities (*fix* and *nif* genes) by horizontal gene transfer of the symbiotic plasmid (35). It is reasonable to assume that the same is true for the denitrification genes as they are interspersed among the *fix* genes on pSymA and both the symbiotic nitrogen fixation genes and the denitrification genes share the same regulator (FixJ). The origin of the denitrification genes may also explain the differences in denitrification efficiency observed between various strains of *S. meliloti*. For example, in a study of 13 *S. meliloti* strains, three failed to denitrify when nitrate was used as a terminal electron acceptor under anaerobic conditions, despite the presence of the appropriate reductases (1). However, when some amount of oxygen was present, all strains tested were able to reduce nitrate to either nitrous oxide or dinitrogen gas (1).

In most bacteria the regulators of the denitrifying pathway (*fixK*, *nosR*, *nnr*) lie in close proximity to the genes that code for the reductase enzymes; this is also true of *S. meliloti*, these regulators are found on the symbiotic plasmid pSymA (33). Other denitrification regulators that have been observed are typically global transcriptional activators found outside the denitrification loci (2). In this study we analyze Adr, a LuxR-like protein located on the chromosome of *S. meliloti*. LuxR family proteins affect a broad number of cell functions that typically act on multiple operons; both FixJ and NarL belong to this superfamily due to the helix-turn-helix homology they share. Adr also appears to play a role in the expression of many cell functions including denitrification, ornithine catabolism (*arc*), sugar metabolism (*smo*), motility, and genes that appear to be involved in respiration (*pnt* and *cyo*) (Supplemental Table S1).

As Adr is a predicted quorum sensing regulator, we examined whether population size or AHLs play a role in how this regulator functions. We observed that the presence or absence of the native AHLs synthesized by *S. meliloti* had no effect on the expression of the denitrification genes. Upon further analysis of the Adr sequence, we found that though this regulator contains a LuxR-like AHL binding domain, key residues for AHL binding are absent. This makes it unlikely that Adr is a traditional AHL-dependent LuxR-like quorum sensing regulator. However, when we examined the expression of the denitrification genes at different growth phases in the presence and absence of Adr, we found that Adr begins to affect expression in between early growth (OD_600_ 0.2) and mid-exponential growth (OD_600_ 0.8). This allows for the possibility that Adr is responding to an environmental or secreted signal that is not traditionally associated with quorum sensing, such as environmental oxygen concentration. In an attempt to induce expression during early growth, we grew cells in spent media that should contain the putative signal molecule, but we were unable to detect any difference in expression between cells grown in fresh media and spent media. However, we do not rule out the possibility of a cell produced molecule triggering the expression of denitrification genes by Adr.

Various factors such as nitrate concentration, oxygen concentration, metal ion availability, and moisture influence denitrification in the laboratory as well as the external environment (2). Denitrification can occur under a range of oxygen concentrations and is no longer considered a strictly anaerobic process. In this study we examined how oxygen influences the expression of the denitrification system in *S. meliloti*. In our genome wide transcriptomic study, we compared the *adr* mutant to wild-type Rm8530. We found that the denitrification pathway is expressed in aerobically grown wild-type cells during stationary phase, but expression is decreased in the absence of *adr*. Previously, Bobik et al. *s*howed that FixJ is the major regulator of denitrification and low oxygen response in *S. meliloti*. When our data is compared side by side with that obtained by Bobik et al., there is a remarkable overlap between the results (17). With this in mind, we examined the relationship between Adr and FixJ. Under our conditions we did not expect the low oxygen dependent regulator of denitrification, FixJ, to be active. However, we observed that when FixJ was removed, the expression of the denitrification genes was reduced 100- to 1000-fold when compared to wild-type levels of transcription. There are several possible explanations for this 1) the oxygen content of our aerobic cultures was low enough to activate FixJ via FixL, 2) unphosphorylated FixJ is sufficient to induce transcription of the denitrification genes, or 3) an unknown factor allows for enhanced aerobic activity of FixJ. Considering the high degree of similarity between the Adr data and the expression data gathered by Bobik, *et al* on FixJ, we consider the third option to be most likely, with the unknown facilitating factor being identified here as Adr. We do acknowledge that oxygen concentrations in stationary phase media are decreased when compared to fresh media. However, it is clear that FixJ and Adr each have an effect on denitrification expression without intentional oxygen removal. Further research is required to determine the mechanism by which the above observed phenomena occur.

Due to the tiered expression control of the denitrification genes, we also included FixK in our study. Under aerobic growth conditions, the presence of both FixJ and Adr, acting through FixK, are required for the optimal expression of the denitrification genes (*nap*, *nir*, *nor*, *nos*) in *S. meliloti*. While the absence of Adr decreases the expression of these genes 5- to 300-fold, removing FixJ has a larger effect on expression. This indicates that the effect of Adr on denitrification is indirect; as proposed above, Adr likely acts to facilitate FixJ-dependent expression when oxygen is present. Since neither FixJ nor Adr affect the other’s transcription, we suspect that a protein-protein interaction may be occurring. Whether Adr functions to stabilize FixJ under free-living oxygenated conditions or help FixJ bind to the DNA is unknown at this time. Further study is required to elucidate the potential interaction occurring between these two proteins.

In addition to transcriptomic studies, we also assessed the ability of Rm8530 and the Rm8530 *adr* mutant to denitrify in a variety of conditions, including different oxygen concentrations and in the presence and absence of nitrate and nitrite. Nitrate represents one of the fixed forms of nitrogen in the environment and an essential component in the biosphere, serving as a nutrient for plants and microorganisms and acting as a final electron acceptor in several bacteria, archaea, and eukaryotes (41). This heavily sought after resource is utilized by microbes involved in performing reactions that contribute to the nitrogen cycle, such as denitrification and dissimilatory nitrate reduction (42). The reduction of nitrate to nitrite is a key reaction in the nitrogen cycle and is the first step of denitrification. Nitrite is found in water and soil environments as a result of the first step of denitrification and as an intermediate of microbial nitrification (43).

Under anaerobic conditions, no growth and no evidence of denitrification was observed in either strain of *S. meliloti*, even after seven days of incubation. Since *S. meliloti* is an aerobe, we surmise that it is not capable of transitioning from respiration to denitrification when there is a complete lack of oxygen. It has also been suggested that most rhizobial denitrification is energy inefficient, since they are observed to grow slowly in comparison to other denitrifiers (44). This observation serves to emphasize the variation in denitrification ability seen between strains of *S. meliloti*. In the previously mentioned study of thirteen *S. meliloti* strains, ten strains were capable of anaerobic growth using nitrate as a terminal electron acceptor (1).

Under aerobic conditions we observed limited denitrification by qualitatively monitoring nitrite levels in the media using nitrite test strips; cells grew normally in both nitrate and nitrite with no survival defects (Figure 6a and 6b), though the *adr* mutant was not as efficient at reducing nitrate and nitrite as the wild-type strain. When cells were grown under microaerobic conditions, a decrease in growth and survival in relation to the aerobic cultures was observed (Figure 6). This growth defect may be explained in several ways. When growing aerobically, *S. meliloti* does not have to rely on denitrification for energy generation. While the process is occurring, it is simply not necessary for the production of proton motive force and ATP because the electrons will pass to oxygen instead of a NO_x_. Once the oxygen concentration in the environment drops, the cells require high oxygen affinity cytochrome oxidases and NO_x_ reductases in order to maintain energy production. Therefore, it is possible that this decrease in growth is due to the lower amount of energy that *S. meliloti* is able to produce under limited oxygen conditions (45).

While nitrite is known to be toxic to *S. meliloti*, we saw no evidence of this during aerobic growth. When the cells were grown microaerobically in the same concentration of nitrite, growth and survival were both reduced (Figure 6c and 6d). In the *adr* mutant, it is reasonable to assume that this reduction in survival is due to the impaired ability of the strain to reduce nitrite, leading to lower energy production and longer exposure to the nitrite. The wild-type strain also has reduced survivability which may be attributable to the closed environment of the growth vial that would allow the accumulation nitric oxide (toxic to *S. meliloti*) before complete reduction to nitrogen gas can occur.

We next explored whether the *nap* and *nir* translation products are functional in *S. meliloti*. Both the nitrate (*nap*) and nitrite (*nir*) reductases in *S. meliloti* are localized in the periplasm. *S. meliloti* only possesses the periplasmic *nap* variant of nitrate reductase; the respiratory nitrate reductase is not present in the genome. Typically, periplasmic nitrate reductases are expressed aerobically in order to remove excess reducing power and provide nitrite for the next step of aerobic denitrification (46). Conversion of nitrite to nitric oxide can be carried out by two types of reductases: the cytochrome *cd1*-type reductase or the copper containing nitrite reductase. *S. meliloti* encodes *nirKV*, a copper containing periplasmic nitrite reductase and an accessory protein which is required for reductase activity (45). The function of both the nitrate and nitrite reductases can be assayed with the addition of the artificial electron donor methyl viologen. When added to whole cells, methyl viologen can donate electrons to enzymes located in the periplasm, but cannot cross the bacterial membrane, allowing for a specific assay of reductases localized in the periplasm (37).

Since the nitrate and nitrite reductases are not oxygen sensitive, we assayed the reductase activities in aerobically grown cells as well as microaerobically grown cells. No denitrifying activity was seen when nitrate was used as a substrate in either growth conditions. This corroborates the results seen in our growth assays. If *S. meliloti* were capable of reducing nitrate to nitrite, we would expect to see the same microaerobic growth defect in cultures with nitrate as those results seen when nitrite was present (Figure 6d). These results lead to the conclusion that the denitrification pathway in Rm8530 is truncated and likely begins at the second step, the reduction of nitrite to nitric oxide. Many denitrifiers are capable of bypassing this low energy reduction of nitrate to nitrite since nitrite is present in the environment (43). For example, similar results were seen in a *napA* gene analysis in *Pseudomonas* isolates; several strains were positive for *nap* gene expression but were incapable of denitrification when nitrate was provided as a substrate (47).

When nitrite was provided as a substrate, we found nitrite reductase activity in both aerobic and microaerobic conditions in wild-type Rm8530. There is limited activity of the nitrite reductase in the *adr* strain under aerobic conditions, supporting our previous data showing a reduction in transcription of these genes in the mutant. Additionally, when the *adr* mutant was grown under microaerobic conditions, there was increased activity of the nitrite reductase compared to aerobic conditions, though levels were not restored to wild-type activity. This increase may be linked to the microaerobic environment. Previous studies have shown that under low oxygen concentrations, FixJ is responsible for *nirK* expression (48). This leads to the possibility that Adr mainly functions when the cells are growing in oxic conditions.

To determine if Adr is necessary for symbiosis with a plant host, we performed symbiosis assays. We found no significant difference between wild-type and the *adr* mutant symbiotic ability, which indicates that Adr is not essential for symbiosis or nitrogen fixation. This suggests that under microoxic conditions, prior to bacteroid differentiation, FixJ in conjunction with FixL are sufficient for mediating expression of the denitrification genes. However, when co-inoculated on the same plant, we observed that the Rm8530 wild-type strain has a competitive advantage in forming nitrogen fixing nodules.

Taken together with our growth and reductase activity results, we propose that a functional Adr, along with FixJ, helps prepare *S. meliloti* for symbiosis during the free-living phase when oxygen concentrations are either too high for FixL to effectively increase phosphorylated FixJ levels, or during the infection stage when oxygen levels fluctuate and begin to decrease as the cells enter the nodule. As seen in Figure 8, we have added Adr to the existing model for denitrification regulation, with a tentative interaction proposed between FixJ and Adr. Whether Adr is influenced by an effector molecule has yet to be elucidated. We observed expression of the denitrification genes in wild-type *S. meliloti* in the presence of oxygen and the absence of denitrification substrates, which can be abolished by the removal of either Adr or FixJ. Aerobic expression of the denitrification genes may serve multiple purposes, such as removal of excessive reducing power or reducing toxic NO_x_ from the environment around the cell. Whether directly or indirectly, Adr influences the aerobic expression of the denitrification genes in *S. meliloti* in conjunction with FixJ. Further study should include elucidation of the possible interaction between FixJ and Adr and the identification and/or isolation of the Adr effector molecule.

## Materials and Methods

### Bacterial strains and media

Strains and plasmids used in this study are listed in Table *S. meliloti* strains were grown at 30° C, 250 rpm in Luria-Bertani media supplemented with 2.5 mM MgSO_4_ and 2.5 mM CaCl_2_ for routine cultures (referred to as LB/MC). *Escherichia coli* cultures were grown in Luria-Bertani media with the appropriate antibiotics at 37° C, 250 rpm. For RNA isolation, *S. meliloti* cultures were grown in tryptone-yeast extract medium supplemented with 3.6 mM calcium chloride (TYC) or minimal low phosphate (19 mM glutamic acid, 55 mM mannitol, 0.1 mM K2HPO4/KH2PO4 mix, 1 mM MgSO_4_, 0.25 mM CaCl_2_, 0.004 mM biotin, pH 7).

To screen for recombinant mutants, strains were grown on minimal glutamate media (MGM) plates (11 g Na_2_HPO_4_, 3 g KH_2_PO_4_, 0.5 g NaCl, 1 g glutamate, 10 g mannitol, 1.5% agar, 1 mg/ml biotin, 27.8 mg CaCl_2_, and 246 mg MgSO_4_); plates were supplemented with 5% sucrose when appropriate. Media for growth curves was supplemented with various nitrogen sources at the following concentrations: 10 mM KNO_3_, or 5 mM NaNO_2_.

Antibiotics were used in the following concentrations when appropriate, streptomycin (Sm) 500 μg/ml, neomycin (Nm) 200 μg/ml, trimethoprim (Tp) 200 μg/ml, spectinomycin (Sp) 100 μg/ml, kanamycin (Km) 25 μg/ml, and chloramphenicol (Cm) 20 μg/ml.

### Construction of the *adr* mutant

*adr* was amplified from *S. meliloti* Rm8530 chromosomal DNA and cloned into the EcoRV site of pPCR-Script, creating the vector p658. The EZ::TN insertion kit (Epicenter) was used to disrupt the cloned *adr* by transposon mutagenesis to create p658Tp. The disrupted *adr* was then cloned into the SpeI site of the suicide vector pJQ200SmSp and the resulting recombinant plasmid (pJQ658Tp) was transformed into DH5α (25). Triparental mating was performed with DH5α pJQ658Tp, MT616 (helper strain), and Rm8530 (49). *S. meliloti* carrying the disrupted copy of *adr* was selected by plating on minimal media supplemented with Tp and 5% sucrose (50). Mutations were confirmed by PCR and phage M12 was used to transduce the *adr* mutation into subsequent strains (51). Primers for mutant construction and confirmation are listed in Table 2.

### Construction of the denitrification mutants

Internal fragments of *fixJ* and *fixK2* were cloned into pK19mobΩHMB creating recombinant vectors harbored in *E. coli* S17-λ*pir* (Table 1). These vectors were provided by Dr. Anke Becker from the Philipps University of Marburg, Germany. The vector carrying *fixK2* was modified by inserting a hygromycin cassette cloned from pMB419 into the Km cassette of the pK19fixK2 backbone. Vectors were transferred via bi-parental mating into *S. meliloti* Rm8530 and recombinants were selected by plating on minimal media with the appropriate antibiotics. Mutations were confirmed by PCR and phage M12 was used to transduce the *fixJ* mutation in Rm8530 *658*::-Tp (51).

To construct the *fixK1 fixK2* double mutant, an internal fragment of *fixK1* was cloned into pVIK112 to create pVIKfixK1. This vector was transferred via tri-parental mating into the Rm8530 *fixK2*::Hy strain. Mutations were confirmed by PCR.

*S. meliloti* Rm2011 containing a Tn5 mutation in *fixL* was also provided by Dr. Anke Becker. This mutation was transferred into Rm8530 by phage M12 and plated on LB agar with the appropriate antibiotics (51). Mutations were confirmed by PCR.

### RNA purification and cDNA synthesis

Bacterial cultures were grown for two days in LB/MC and appropriate antibiotics. A 1:100 dilution was used to inoculate 20 ml of TY media supplemented with Sm. Cultures were grown aerobically to OD_600_ 0.2 (lag), 0.8 (mid log), or 1.2 (stationary). After reaching the appropriate growth stage, 1.5 ml aliquots of culture were harvested by centrifugation (14,500 rpm for 2 minutes at 4° C), immediately frozen in liquid nitrogen, and stored at -80° C for future use. RNA purification was performed using the RNeasy Mini Kit (Qiagen) with slight modifications. Briefly, cells were thawed on ice then resuspended in 10 mM Tris HCl (pH 8) and RLT buffer provided from the Qiagen kit (supplemented with β-mercaptoethanol). The cells were transferred to FastProtein tubes (Qbiogene) and disrupted using an MP FastPrep-24 ribolyser (40s, speed 6.5). Spin column purification was performed according to the RNeasy Mini Kit RNA purification protocol. After the first round of purification, samples were treated with Qiagen on column RNase-free DNase. The RNA samples were eluted and DNase treated a second time with the Ambion TURBO RNase-free DNase which was followed by an RNA clean up step. Concentration of RNA was determined by Nanodrop and DNA contamination was assessed by qRT-PCR. cDNA for each strain was synthesized with the Ambion RETROscript kit according to the manufacturer's protocol. 1 μg of total RNA was used per cDNA synthesis reaction.

### Affymetrix GeneChip hybridization and expression analysis

The cDNA synthesis was performed using 10 μg of RNA harvested from cells grown to OD_600_ 1.2. Hybridization of the cDNA to the GeneChip Medicago Genome Array (Affymetrix, Santa Carla, CA) was performed at the Core Microarray facility at UT Southwestern Medical Center (Dallas, TX) as previously described (27, 32). The GeneChip Scanner 3000 was used to measure the signal intensity of each array. Affymetrix GeneChip Operating Software, (GCOS v 1.4) was used to generate the .CEL files. Comparative analysis of the control and experimental expression were represented in terms of M-value (signal log ratio) which also indicated an increase, decrease, or lack of change in expression of a gene in the mutant with respect to the wild-type. An M-value ≥ 1 (2-fold change) with a *p*-value of ≤ 0.05 were considered significant.

### Quantitative real-time PCR

Oligonucleotide sequences used for qRT-PCR are listed in Table 3. The reaction mixture for qRT-PCR analysis contained 0.3 μM of sense primer, 0.3 μM of antisense primer, 0.5X of SYBR green 1 (Sigma), 0.5 Omni Mix HS PCR bead (contains 1.5 U *Taq* DNA polymerase, 10 mM Tris-HCl [pH 9.0], 50 mM KCl, 1.5 mM MgCl2, 200 μM deoxynucleotide triphosphate, and stabilizers), and 1 μl of cDNA. Total reaction volume was 25 μl. Analysis was performed using a Cepheid Smart Cycler, version 2.0c as previously described (32). Expression analyses were conducted in triplicate. The expression of *SMc00128* was used as an internal control and for normalization, as described previously (52, 53). Expression analysis were performed as three independent experiments.

### Growth analysis

Cells for all growth curves were grown to saturation in LB/MC with appropriate antibiotics. Aerobic growth curves were performed using a Tecan plate reader (29° C, shaking at 250 rpm). Cells were diluted 1:100 in TYC media, TYC with 10 mM KNO_3_, or TYC with 5 mM NaNO_2_ and added to a 96 well plate. OD_600_ readings were taken every 30 minutes.

Microaerobic conditions were also tested. Starter cultures were diluted to an initial OD_600_ of 0.2, then 2 ml of each culture was added to each vial. Once sealed, the vials were sparged with a mixture of 2% oxygen, 98% argon for one minute. Vials were incubated at 30° C, shaking at 250 rpm and OD_600_ was measured every 24 hours. We refer to aerated cultures to differentiate between aerobic cultures grown with free gas exchange with the environment and intentionally oxygen restricted (microaerobic) cultures.

Viable counts were performed for both aerobic and microaerobic cultures. Several dilutions were plated on LB agar with the appropriate antibiotics. Plates were incubated for several days at 30° C before colonies were counted by hand. Nitrite levels of cultures were qualitatively assessed using nitrite test strips (0-80 mg/L or 0-10 mg/L, EMD Millipore).

### Media complementation assay

Cells were grown in the same manner as those grown for RNA pellets. Cultures were grown to OD_600_ of 1.2 in TYC media with Sm. Cells were removed from the media by centrifugation (3 × 6000 rpm, 30 minutes, 4° C) and the resulting supernatant was passed through a 0.22 μm filter (Fisher) and stored at 4° C. Glucose (100 mM) and glutamate (19 mM) were added to the spent media to replace depleted nutrients. To determine if the spent media contained effector molecules that are detected by or that activate *adr*, cultures were grown to OD_600_ of 0.2 and then harvested for RNA in the manner described above.

### Methyl viologen assay for nitrate and nitrite reductase activity

Starter cultures were grown aerobically in TYC media with appropriate drugs for two days. Cells were then spun down (2 minutes, 14,000 rpm), and diluted to an initial OD_600_ of 0.2 in MLP supplemented with Sm and 0.5 μM NaMoO_4_. Cells grown aerobically were harvested at OD_600_ of 1.2 by spinning down 1 ml of culture (two minutes, 14,000 rpm, 4° C), removing the supernatant, and immediately freezing the cell pellet in liquid nitrogen. Microaerobically grown cells were harvested at OD_600_ of 0.5-0.6 (stationary phase for microaerobic cultures).

Samples were assayed for either nitrate or nitrite reductase activity using methyl viologen as the electron donor as previously described (37, 54). Cell pellets were resuspended in 10 mM Hepes buffer (pH 7) and 1 mM methyl viologen to a final volume of 1 ml. The cuvettes were sealed and sparged for five minutes with nitrogen gas. Methyl viologen was reduced by adding aliquots of freshly prepared aqueous sodium dithionite until a steady state absorbance of 1-1.5 at 600 nm was obtained. Substrate (10 mM nitrate or nitrite) was added, and the rate of absorbance decrease measured. The rate of substrate-dependent methyl viologen oxidation was calculated using 13 mM-1 cm-1 as the extinction coefficient of reduced methyl viologen (55). Protein concentrations were measured by the Bradford method (Bio-Rad) and used to determine specific activities. All strains were measured in triplicate.

### Plant symbiosis assays

Infection assays of *Medicago sativa* were performed to determine nodulation and nitrogen fixation efficiency of the *S. meliloti* mutant strains Rm8530 *adr*, Rm8530 *fixJ*, and Rm8530 *fixJ adr* compared to wild-type Rm8530. *M. sativa* was inoculated with *S. meliloti* strains on Jensen’s agar plates as previously described (56). Plants were grown in a 16-hour light cycle at 20° C and in 65% humidity. Weekly inspections of roots and plant health were performed beginning the second week post inoculation. Nitrogen fixing nodules, empty nodules, and plant height were recorded for approximately 60 plants per strain tested. Data shown was collected fourth week post inoculation.

### Plant symbiosis competition assay

Five dilutions (100:1, 10:1, 1:1, 1:10, and 1:100) of Rm8530 to Rm8530 *adr* were tested for competitive nodulation of *M. sativa.* Strains were grown for two days in LB/MC with antibiotics, washed three times with sterile water, then diluted 1:100 in water. These dilutions were then mixed to obtain the appropriate ratio of wild-type to mutant. A portion of the mixed inoculum was diluted and plated on the appropriate antibiotics to determine cell viability. *M. sativa* seedlings were inoculated with 1 ml of the dilution and plants were grown in the same conditions discussed above. Harvested nodules were washed in 50% bleach for five minutes and rinsed three times with sterile water. After rinsing, each nodule was added to a well of a microtiter plate and crushed in LB/MC supplemented with 0.3 M glucose.

The bacteria released from the crushed nodules were diluted, divided in half, and plated on LB/MC Sm and LB/MC Tp. Colony PCR was used to confirm the genetic background of the strains. Nodulation competitiveness was tested twice for each dilution.

## Acknowledgements

We thank Anke Becker and Jacques Batut for providing strains. We also thank Stephen Spiro for his assistance with the methyl viologen and nitrite detection assays and for reviewing this manuscript. Finally, we thank Nymisha Avadhanam for reading and giving feedback on the manuscript. Our work was supported by National Science Foundation grant MCB-9733532 and National Institutes of Health grant 1R01GM069925 to J.E.G.

